# Defective intestinal repair after short-term high fat diet due to loss of efferocytosis

**DOI:** 10.1101/790220

**Authors:** Andrea A. Hill, Myunghoo Kim, Daniel F. Zegarra-Ruiz, Hyo Won Song, Michael C. Renfroe, Angela M. Major, Wan-Jung Wu, Kendra Norwood, Gretchen E. Diehl

**Affiliations:** Alkek Center for Metagenomics and Microbiome Research and the Department of Molecular Virology and Microbiology, Baylor College of Medicine, Houston TX 77030, USA; Department of Animal Science, Pusan Nation University 50463 Republic of Korea; Texas Children’s Pathology, Division of Anatomic Pathology, Texas Children’s Hospital, Houston TX 77030, USA; Biology of Inflammation Center, Baylor College of Medicine, Houston TX 77030, USA

## Abstract

Defective tissue repair is a hallmark of many inflammatory disorders including, inflammatory bowel disease (IBD). While it is clear that high fat diets (HFD) can exacerbate inflammatory disease by increasing inflammation, the direct effect of lipids on tissue homeostasis and repair remains undefined. We show here that short term exposure to HFD directly impairs barrier repair after intestinal epithelial damage by interfering with recognition and uptake of apoptotic neutrophils by intestinal macrophages. Apoptotic neutrophil uptake induces macrophage IL-10 production, which is lacking after intestinal damage in the context of HFD. Overexpression of IL-10 rescues repair defects after HFD treatment, but not if epithelial cells lack the IL-10 receptor, highlighting the key role of IL-10 in barrier repair. These findings demonstrate a previously unidentified mechanism by which dietary lipids directly interfere with homeostatic processes required to maintain tissue integrity.

## Introduction

Within the intestine, a single layer of epithelial cells separates our internal tissues from luminal contents. Under normal conditions, this forms a selective barrier, allowing passage of nutrients and excluding harmful substances and intestinal microbes. Injury to this barrier increases entry of luminal contents into the tissue. Unresolved barrier damage underlies many chronic inflammatory diseases including inflammatory bowel disease (IBD).

During infection or trauma, a pro-inflammatory immune response is critical to contain and clear any infectious agents as well as prime the tissue for repair^1^. First, tissues and tissue resident immune cells detect the damage and recruit circulating neutrophils. Neutrophils extravasate into the tissue to clear any invading organisms. Neutrophil effector mechanisms can damage tissue environments and so must be rapidly contained. One mechanism is the induction of programmed cell death in infiltrating neutrophils. This cell death is also an important cue in resolution of inflammation^1^. Apoptotic neutrophils release mediators that both limit neutrophil recruitment to the tissue and also act as bridging molecules to stimulate uptake by macrophages, also known as efferocytosis. Many studies highlight the role of macrophage efferocytosis, particularly clearance of neutrophils, as a critical cue in macrophage switch from an anti-microbial, pro-inflammatory response to a pro-tissue repair, anti-inflammatory responses including production of IL-10 and TGFβ^2,3^. This further dampens recruitment of neutrophils into the tissues and is critical for a return to homeostasis. Defects in macrophage efferocytosis and subsequent impaired pro-repair responses has been linked to many chronic inflammatory diseases including atherosclerosis, cardiovascular disease, COPD and autoimmune disease, including type 1 diabetes and lupus^4^.

The intestinal tract is exposed to a variety of environmental factors with the potential to disrupt tissue homeostasis, many of which may contribute to disease pathology. One factor is the consumption of dietary lipids and increased intake of diets high in fat (HFD) is a risk factor for chronic inflammatory diseases, including IBD^5^. It is well established that long term exposure to HFD can suppress macrophage anti-inflammatory functions and promote pro-inflammatory responses, supporting local and systemic inflammation^6-8^. However, whether dietary lipids directly impair anti-inflammatory intestinal homeostatic functions and tissue repair is incompletely understood.

Here we show that, after intestinal damage, short term HFD exposure impairs intestinal barrier repair. We find, after intestinal damage, dietary lipids directly interfere with macrophage recognition and uptake of apoptotic cells and subsequent IL-10 production. This is due to lipids blocking interactions between apoptotic cells and the efferocytosis receptor CD36, which also binds dietary lipids. This results in increased apoptotic neutrophil accumulation in the tissue. Overexpression of IL-10 rescues repair defects after HFD, but not if epithelial cells lack the IL-10 receptor, highlighting the key role of IL-10 in barrier repair. These findings demonstrate a previously unidentified mechanism by which dietary lipids, a risk factor for intestinal disease, can directly interfere with homeostatic processes required to maintain tissue integrity.

## Results

### Delayed epithelial repair after intestinal injury in short term HFD fed mice

To understand the impact of high fat diet (HFD) on intestinal barrier repair, we treated mice with high or low fat diet (LFD) for one week before exposure to dextran sodium sulfate (DSS) for 5 days or a single dose of doxorubicin (DOX), both of which cause intestinal epithelial damage^9,10^. Mice remained on their respective diets throughout the experiment. Without DSS exposure, mice in both groups had comparable body weight with normal glucose and insulin tolerance (Fig. 1a, Supplementary Fig. 1a, b). This lack of metabolic changes allowed us to assess the direct impact of HFD on intestinal damage prior to systemic inflammation caused by long-term HFD exposure. After DSS treatment, mice exposed to HFD or LFD initially lost a comparable amount of weight. While LFD mice recovered initial weight, HFD mice showed sustained weight loss (Fig. 1a). We used histopathological analysis to assess whether HFD resulted in increased tissue damage and inflammation. In both HFD and LFD treated animals we found pathology in both distal colon and cecum after DSS treatment as previously described^11^. At day 5 of DSS treatment, we found similar tissue pathology and myeloid cell infiltration in both HFD and LFD mice (Fig. 1b, c, Supplementary Fig 2a, b), indicating that diet was not amplifying initial damage or inflammation as is seen after long term exposure to HFD^12-14^. In LFD treated mice, damage was partially resolved as we see reduced tissue infiltration of immune cells and increased restitution of the epithelial barrier by day 9 post DSS treatment. In contrast, in HFD treated mice, we observed prolonged tissue damage and cellular infiltration (Fig. 1b, c). As with DSS, after DOX treatment, HFD mice showed extended weight loss as compared to LFD mice (Supplementary Fig. 3a).

**Figure 1.**
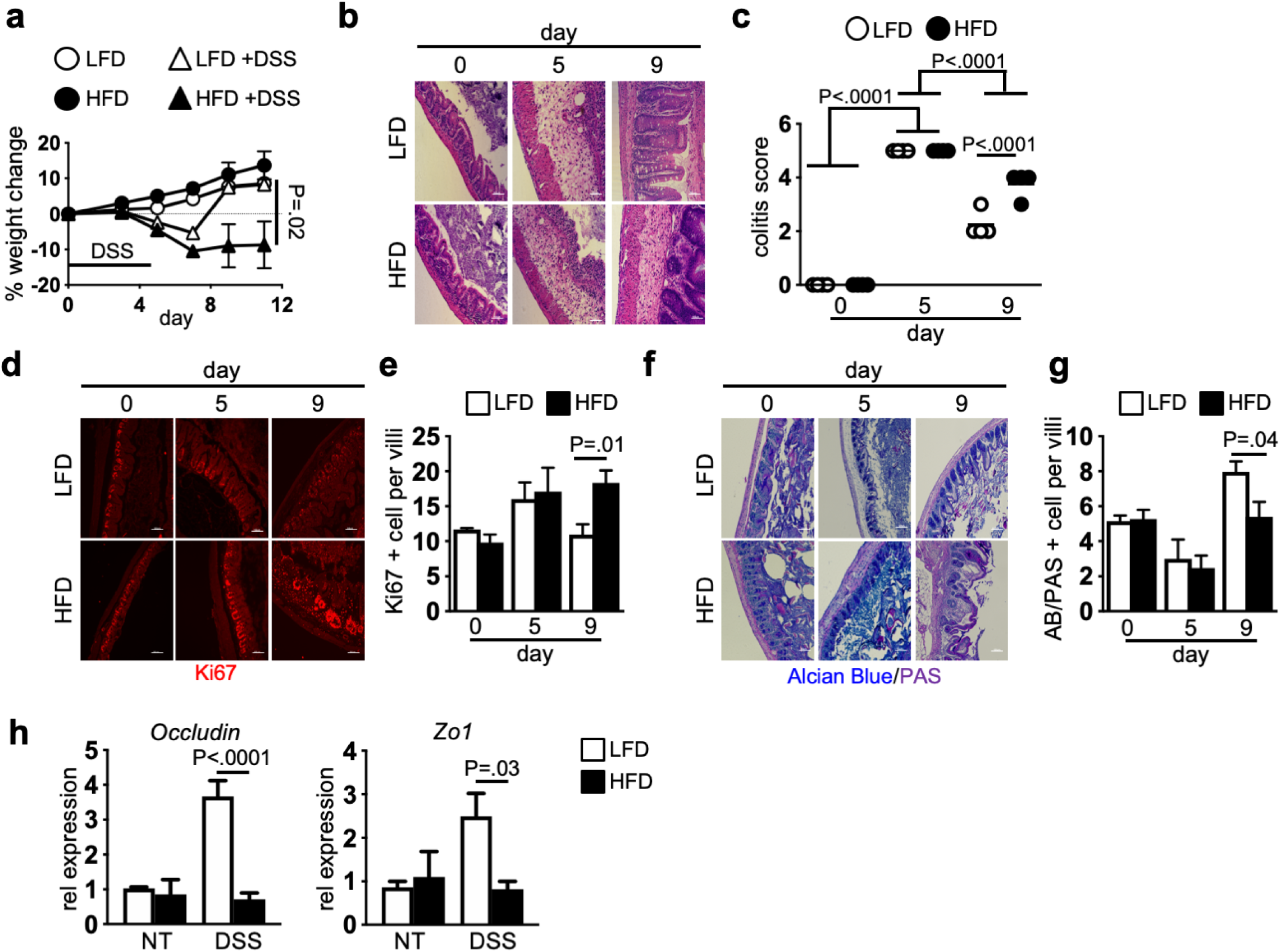
Reduced resolution of intestinal damage after short term high fat diet. WT mice were fed LFD or HFD for one week. Mice were then left untreated or treated with 2% DSS in the drinking water for 5 days. (**a**) Body weight (n=15 mice per group), (**b**) representative H&E stained cecum and (**c**) blinded colitis score at indicated day post DSS treatment. (**d**) Representative image and (**e**) quantification of proliferating cells (Ki67, red) at indicated day post DSS treatment. (**f**) Representative image and (**g**) quantification of Alcian Blue/PAS staining for goblet cells (dark blue) in cecum. n= 5 mice per group, average of 3 HPF images per mouse (**c, e, g**). Scale bar equals 100 microns. (**h**) Occludin and zo-1 gene expression in cecum of LFD and HFD mice after DSS treatment. n=4 mice per group. Data are presented as mean ± SEM. Statistical significance was analyzed by Student’s T test (**a, e, g, h**) or One-way ANOVA with Tukey’s posttest (**c**).

To determine if HFD increased susceptibility of mice to additional models of intestinal damage, we infected LFD and HFD treated mice with *Citrobacter rodentium*, a model of infectious colitis. We did not observe increased weight loss in HFD mice and both HFD and LFD mice had similar histopathology scores (Supplementary Fig. 3b-c). These findings demonstrate that while HFD does not increase susceptibility to intestinal damage or infectious challenge, HFD feeding extends pathology and delays recovery from intestinal damage.

Changes in microbiota composition is associated with increased inflammation and can support inflammation after DSS treatment^9^. Altered microbiota composition is suggested to be a mechanism of HFD driven intestinal inflammation^15^. To determine if microbiota changes after HFD were driving increased pathology, we performed microbial composition analyses by 16S rRNA sequencing. We found modest compositional differences in the intestinal microbiota between the groups after one week of dietary treatment (Supplementary Fig. 4a, c). By day 5 of DSS challenge, there was no difference in microbiota composition between treatment groups (Supplementary Fig. 4b, d). This indicates that while diet can alter microbiota composition, inflammatory insult and damage is a stronger driver, with both groups exposed to similar microbes during resolution of damage.

One of the first steps in repairing intestinal epithelial damage is proliferation of the intestinal stem cell compartment followed by differentiation to repopulate lost cell types^16^. We found similar epithelial proliferation between HFD and LFD fed mice before DSS treatment. Early after DSS treatment, we observed equivalent increased proliferation in both groups (Fig. 1d, e). While proliferation decreased over time in the LFD mice, it remained elevated in the HFD mice (Fig. 1d, e, Supplementary Fig. 5). Along with increased proliferation we also found decreased repopulation of goblet cells and mucus in HFD mice as compared to LFD mice (Fig. 1f, g, Supplementary Fig. 5). In order to repair intestinal damage, tight junctions between epithelial cells must be reestablished. This paracellular barrier prevents microbial translocation into the tissue^17^. Expression of tight junction proteins occludin and zonula occluden-1 (zo-1), which together are the major tight junction proteins^18^, did not differ between LFD and HFD mice before DSS treatment (Fig. 1h). After DSS treatment, we found upregulated expression of occludin and zo1 in LFD but not HFD treated mice (Fig. 1h). Together, these findings demonstrate that overall intestinal epithelial repair processes are impaired in HFD after intestinal injury.

### Increased neutrophil accumulation limits damage repair after DSS in HFD treated mice

A marker of pathology after intestinal damage is increased numbers of apoptotic or necrotic cells within the tissue. Further, accumulation of apoptotic cells and cellular debris within the tissue after injury is associated with defective repair^19^. Using immunofluorescence (IF) staining for TUNEL which detects double strand breaks as found in apoptotic cells, we found that HFD alone did not result in increased numbers of apoptotic cells in the intestine (Fig. 2a, b). In early time points after DSS treatment, apoptotic cell numbers are equivalent in both HFD and LFD mice. While these numbers did not increase over time in LFD treated mice, in HFD treated mice, we found increased apoptotic cells within intestinal tissue and lumen (Fig. 2a, b).

During intestinal damage monocyte-derived macrophages and neutrophils are recruited to sites of injury and their numbers decrease after resolution of damage. Increased presence of inflammatory macrophages, monocytes and neutrophils in intestinal tissue supports inflammation^20,21^. We used immunofluorescence staining and flow cytometric analysis to assess monocyte, macrophage and neutrophil recruitment in LFD and HFD mice after intestinal injury. DSS treatment resulted in increased monocytes, macrophages, and neutrophils in the tissue (Fig. 2c, d and Supplementary Fig. 2, 6). We found no difference in macrophage numbers between HFD and LFD treated mice (Supplementary Fig. 6d). In contrast, while we found equivalent neutrophil numbers in intestinal tissue of HFD and LFD treated mice at day 5 after DSS treatment (Fig. 2d and Supplementary Fig. 7b), at later timepoints, they further increased in HFD fed mice (Fig 2c, d). Neutrophils are short lived cells which die by apoptosis to prevent release of cytotoxic intracellular components^1^. If not cleared, neutrophils can undergo secondary necrosis which can amplify tissue pathology and limit intestinal barrier repair^22^. By immunofluorescence analysis, we find the majority of the TUNEL positive cells within the tissue of HFD and LFD treated mice after DSS challenge are neutrophils with TUNEL positive neutrophils increasing at later stages after intestinal injury after HFD treatment (Fig. 2c and e, Supplementary Fig. 6 and 7). We also find increased expression of neutrophil chemoattractant proteins, CXCL1 and CXCL2, in HFD fed compared to LFD fed mice after injury (Fig. 2f), indicating continued neutrophil recruitment and reduced resolution of tissue injury in the presence of HFD.

**Figure 2.**
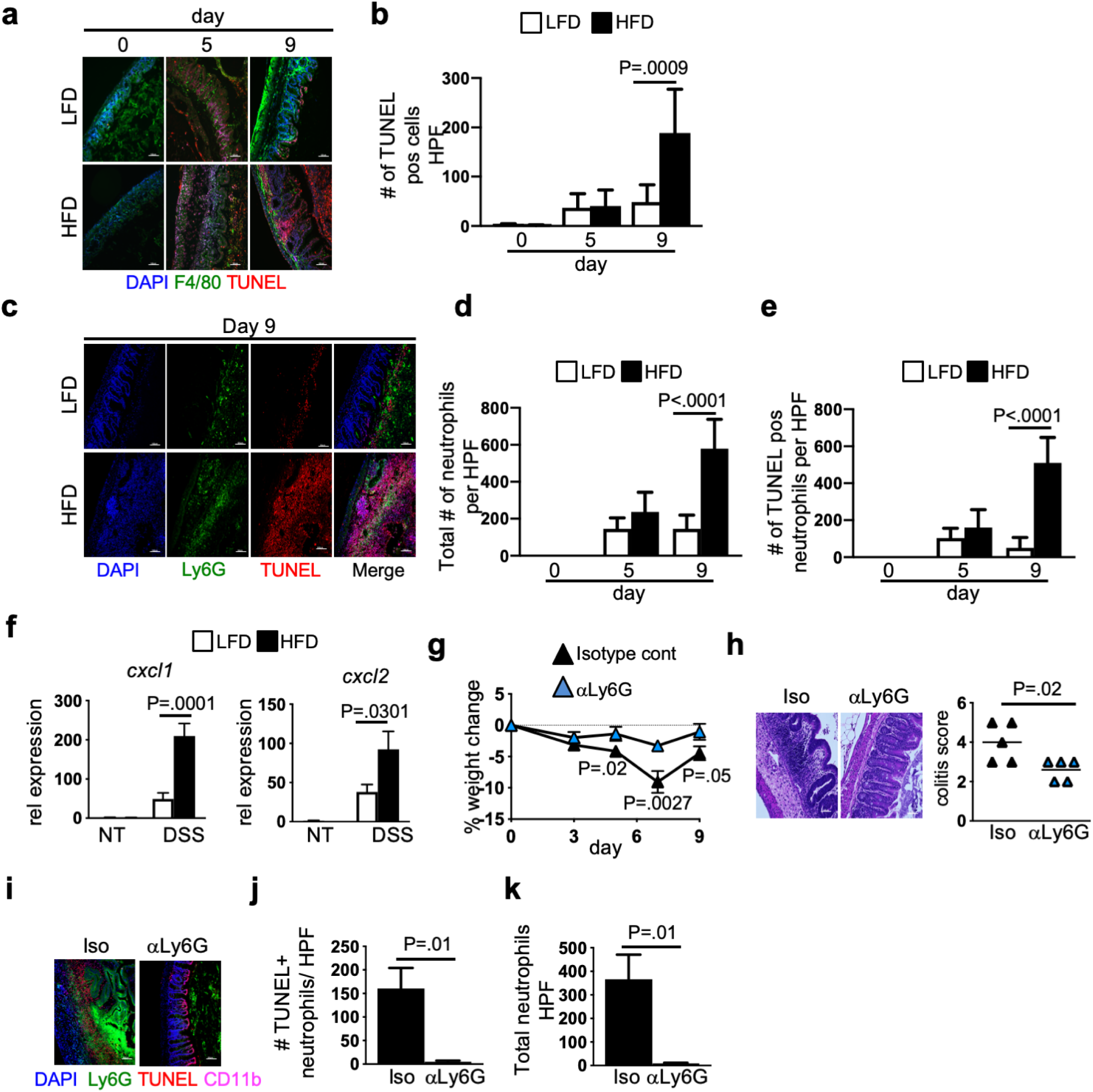
Increased apoptotic neutrophils in intestines of HFD mice after damage. (**a**) Representative image stained for F4/80 (green), nuclei (DAPI, blue), apoptotic cells (TUNEL) CD11b (magenta) and (**b**) quantification of apoptotic cells (TUNEL^+^) in LFD and HFD control and DSS treated mice (n=5). (**c**) Representative image for macrophages (F4/80^+^), nuclei (DAPI), apoptotic cells (TUNEL^+^), neutrophils (Ly6G^+^) in LFD and HFD control and day 9 after DSS treatment (n=5). (**d**) Quantification of total number of neutrophils and (**e**) TUNEL^+^ neutrophils in (**c**) per high-powered field (n=5). (**f**) *Cxcl1* and *cxcl2* gene expression in cecum of LFD and HFD mice after DSS treatment. n=5 mice per group. (**g**) Body weight of HFD DSS mice treated with anti-IgG2a control or anti-Ly6G antibody starting at day 4 post DSS (n= 20). (**h**) Representative H&E image and blinded colitis score for cecum from isotype control and neutrophil depleted HFD DSS treated mice (n= 5). (**i**) Representative immunofluorescence image stained as in (**f**) of neutrophil depleted HFD DSS treated mice. (**j**) Quantification of TUNEL positive neutrophils (Ly6G^+^) in panel (**i**) per high-powered field. (**k**) Quantification of total number of neutrophils in panel (**i**) (n= 5, average counts of 3 HPF images per mouse per “n”). Scale bar equals 100 microns. Data are presented as mean ± SEM. *p<0.05. Statistical significance was analyzed using One-way Anova with Tukey’s posttest (**b, d, e**) or Student’s T test (**f, g, h, j, k**).

Increased neutrophil numbers and inefficient clearance of apoptotic neutrophils can result in impaired tissue resolution^21,22^. To investigate whether accumulation of neutrophils was required for increased pathology after HFD, we depleted neutrophils in HFD mice after the start of DSS treatment. Neutrophil depletion in HFD DSS treated mice resulted in improved body weight as compared to isotype control antibody treated animals (Fig. 2g) with improved histopathology, decreased epithelial proliferation, and restored goblet cell numbers as compared to isotype control antibody treatment (Fig. 2h and Supplementary Fig. 8a, b). This was accompanied by decreased numbers of total neutrophils and TUNEL positive cells within the tissue (Fig. 2i-k). These results demonstrate that neutrophils are required for impaired intestinal barrier repair in HFD fed DSS treated mice.

### HFD treatment impairs barrier repair by limiting macrophage production of IL-10 after intestinal injury

Phagocytosis of apoptotic cells leads to macrophage expression of a number of anti-inflammatory and tissue repair factors such as IL-10^23^. IL-10 both supports epithelial barrier repair^24,25^ and increases macrophage efferocytosis capacity^26^. While we find IL-10 is strongly upregulated in intestinal tissue of LFD fed mice after DSS treatment, IL-10 is not upregulated in intestines of DSS treated mice exposed to HFD (Fig. 3a). In the intestine, we and others have found macrophages are the major source of IL-10^27-29^. After DSS exposure, IL-10 expression by macrophages from mice on HFD is reduced as compared to macrophages from mice on LFD (Fig. 3b). Further, total tissue and macrophage expression of pro-inflammatory genes such as TNFα were not altered after HFD treatment (Supplementary Fig. 9a, b). These findings suggest HFD limits macrophage IL-10 response to tissue damage without increasing inflammatory functions.

**Figure 3.**
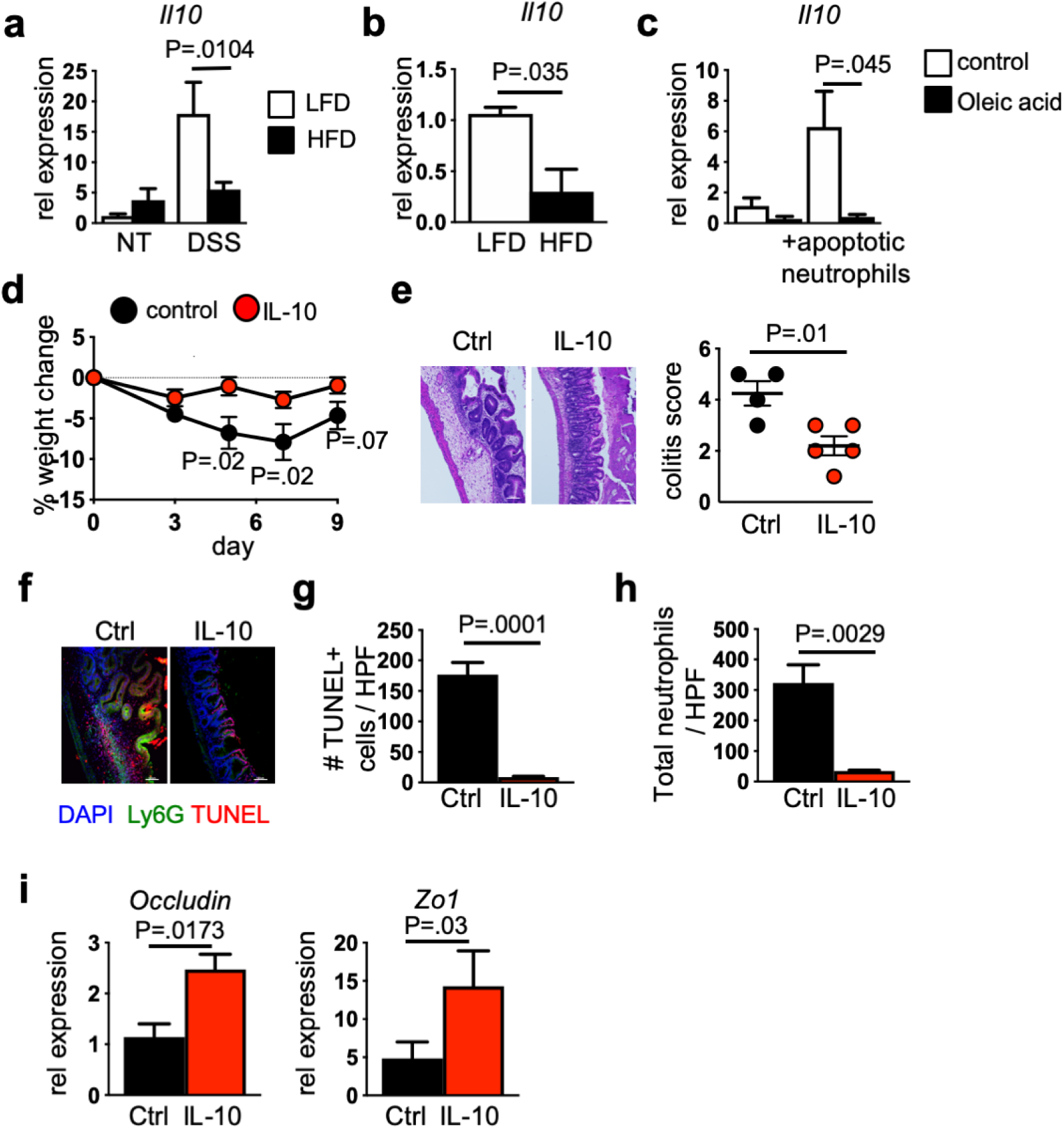
IL-10 rescues intestinal repair defect in HFD mice. (**a**) *IL-10* expression from cecum of LFD and HFD control and DSS treated mice (n=6 LFD, n=6 HFD, n=7 LFD DSS, n=10 HFD DSS mice). (**b**) Gene expression of IL-10 in sorted intestinal macrophages from LFD and HFD DSS mice (n=5 per group, pooled samples from 3 mice per “n”). (**c**) Gene expression of IL-10 in control and oleic acid pre-treated macrophages after apoptotic neutrophil exposure (n= 4 replicate experiments). (**d**) Body weight change in HFD DSS mice after hydrodynamic gene delivery of an IL-10–producing plasmid or control (n= 8 per group). (**e**) Representative H&E staining and blinded colitis scores (n=5 per group) of cecum from mice in panel (**d**). (**f**) Representative immunofluorescence staining for macrophages as in Figure 2 of HFD DSS mice with hydrodynamic gene delivery of an IL-10–producing or control plasmid. (**g**) Quantification of TUNEL positive cells per high-powered field (n=5 mice per group). (**h**) Quantification of total neutrophils in panel (**f**). (**i**) Occludin and zo1 expression in cecum of mice in (**d**) (n=8 mice per group). Data are presented as mean ± SEM. Statistical significance was analyzed by One-way Anova with Tukey’s posttest (**a**) and Student t Test (**b, c, d, e, g, h, i**).

In a number of inflammatory disorders, lipids, including those derived from the diet, can inhibit normal macrophage tissue repair functions and sustain inflammation. For example, in obese adipose tissue, macrophages accumulate lipids which promotes secretion of inflammatory mediators such as TNFα^30^ and decreases expression of IL-10^7^. To understand if intestinal macrophages accumulated intracellular lipids as found in obesity, we performed Oil-red-O staining of sorted intestinal macrophages from LFD and HFD mice and found no differences in lipid accumulation (Supplementary Fig. 10). This suggests lipid accumulation does not contribute to loss of intestinal macrophage IL-10 in HFD treated mice after intestinal injury.

To determine if dietary lipids directly alter IL-10 expression in macrophages in response to apoptotic cells we pretreated bone marrow derived macrophages (BMDMs) with oleic acid, a dietary lipid found in the HFD, before exposing them to apoptotic neutrophils. Oleic acid treatment did not result in increased macrophage apoptosis (Supplementary Fig. 11). Apoptotic neutrophil exposure resulted in macrophage IL-10 production. However, IL-10 was not upregulated after lipid pretreatment (Fig. 3c). Overall, these findings demonstrate that dietary lipids directly impair macrophage IL-10 production in response to apoptotic neutrophils.

To understand if IL-10 was sufficient to protect mice from intestinal damage in the presence of HFD, we overexpressed IL-10 in vivo and found it was sufficient to protect HFD treated mice from increased weight loss after DSS treatment (Fig. 3d). We also observed improved tissue histology, reduced numbers of apoptotic cells in the tissue, and restored goblet cell formation alongside reduced neutrophil numbers and epithelial proliferation (Fig. 3e-h and Supplementary Fig. 12a, b). Further, overexpression of IL-10 increased expression of gap junction proteins occludin and zo-1 (Fig. 3i). These finding demonstrate IL-10 is sufficient to normalize barrier repair in HFD treated mice after intestinal injury.

### Epithelial cell IL-10 signaling is required for tissue repair after DSS

To identify the target of macrophage IL-10 production, we overexpressed IL-10 in HFD mice lacking IL-10 receptorα (IL-10Rα) on macrophages or epithelial cells. IL-10Rα on macrophages is dispensable as IL-10 overexpression rescues HFD treated mice lacking macrophages IL-10Rα after DSS treatment (Supplementary Fig.13). In contrast, IL-10 overexpression is unable to rescue bodyweight, colitis score or apoptotic neutrophil accumulation in DSS and HFD treated mice where epithelial cells lack IL-10Rα (Fig. 4a-e). These data demonstrate that IL-10 signaling on intestinal epithelial cells is required for resolution of barrier injury.

**Figure 4.**
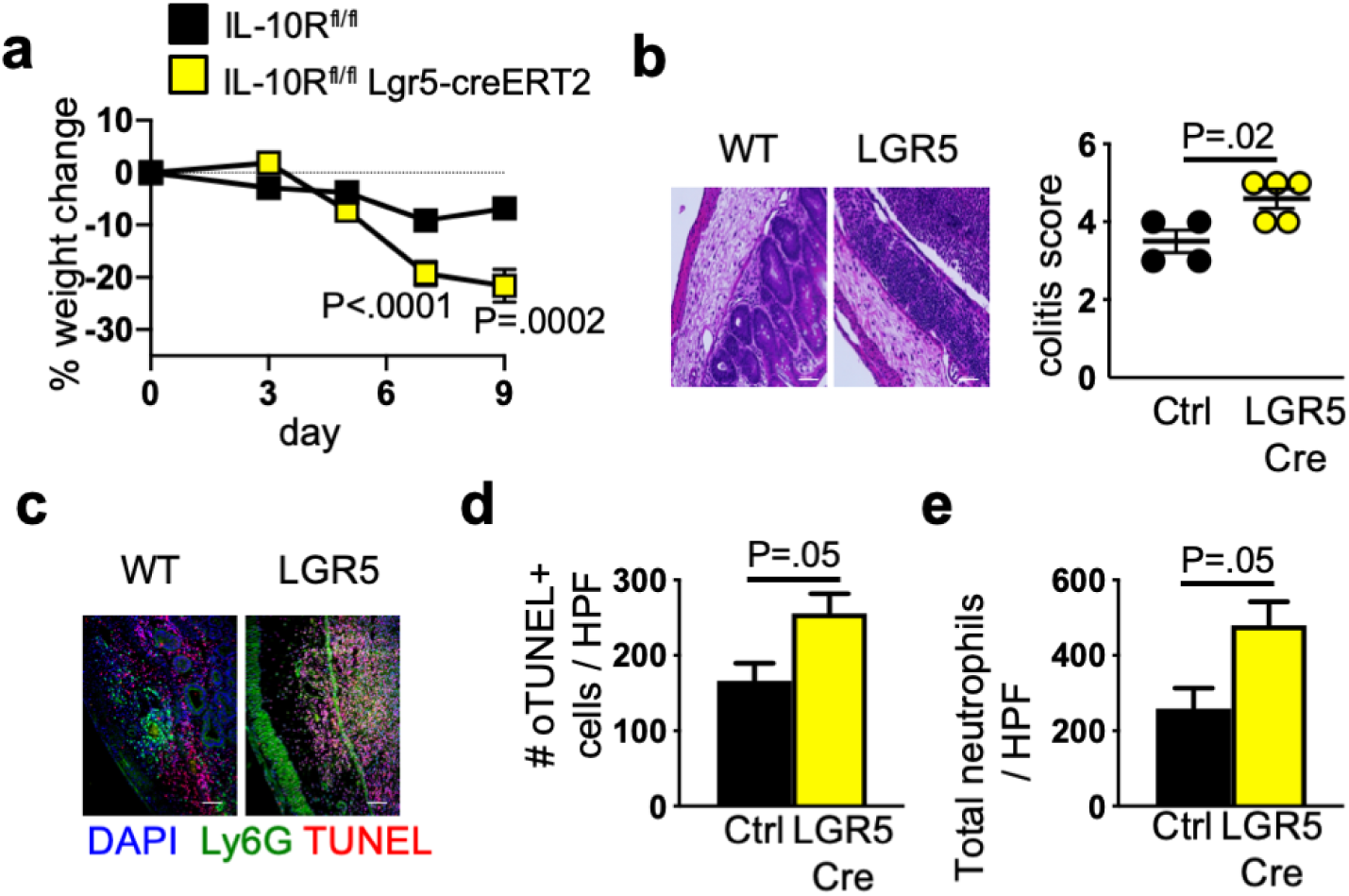
Loss of IL-10 signaling in epithelial cells inhibits intestinal repair in HFD mice. (**a**) Weight loss in tamoxifen treated IL-10R^fl/fl^ (control) and IL-10R^fl/fl^ LGR5-creERT2 mice treated with HFD and DSS and hydrodynamic delivery of an IL-10–producing plasmid (n= 11 no cre n=7 cre). (**b**) Representative H&E image and blinded colitis score from cecum of mice in panel (**a**) (n=4 IL-10R^fl/fl^ and n=5 IL-10R^fl/fl^ LGR5-creERT2). **(c)** Representative immunofluorescence staining for macrophages as in Figure 2 of HFD treated control and IL-10R^fl/fl^ LGR5-creERT2 mice exposed to DSS. (**d**) Quantification of TUNEL positive neutrophils (Ly6G^+^) in panel (**c**) per high-powered field. (**e**) Quantification of total number of neutrophils in panel (**c**) (n= 5, average counts of 3 HPF images per mouse per “n”). Scale bar equals 100 microns. Data are presented as mean ± SEM. Statistical significance was analyzed by Student t Test (**a, b, d, e**).

### HFD exposure decreases efferocytosis by intestinal macrophages

Efferocytosis is a critical function of tissue macrophages^4^. To understand if macrophages from DSS treated mice exposed to HFD were defective in uptake of apoptotic cells, we exposed flow sorted intestinal macrophages to TAMRA labeled apoptotic neutrophils. We find reduced efferocytosis of apoptotic neutrophils in sorted macrophages from mice with intestinal injury after HFD (Fig. 5a).

**Figure 5.**
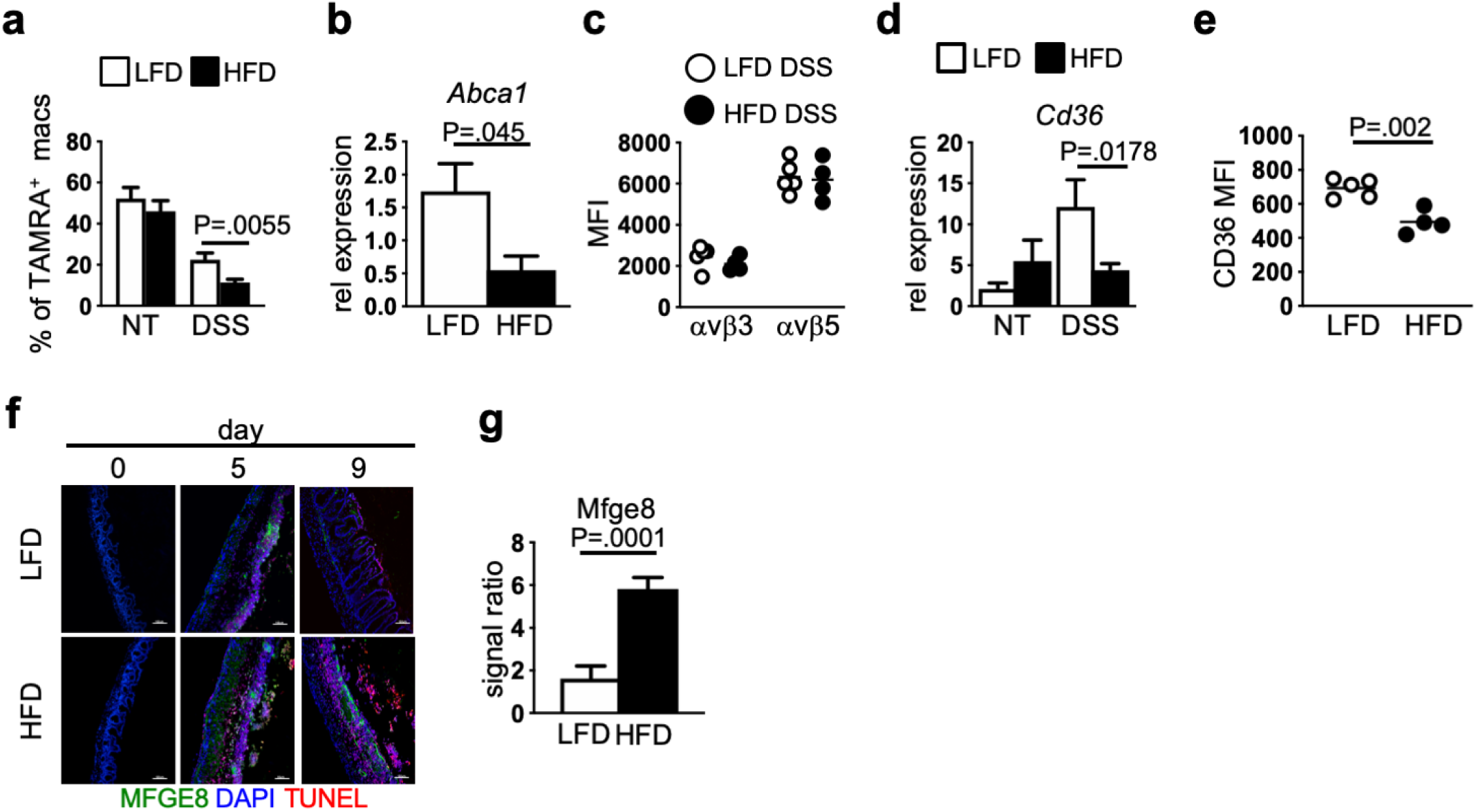
Dietary lipids directly inhibit macrophage efferocytosis. (**a**) Quantification of percent of tethered TAMRA positive neutrophils in sorted macrophage from LFD and HFD mice after DSS treatment (LFD DSS (n=4), HFD DSS (n=4)) (**b**) Gene expression of *Abca1* in sorted intestinal macrophages (n= 4, 3 mice per group pooled per “n”). (**c**) Flow cytometry analysis for surface expression of αvβ3/5 in intestinal macrophages of LFD and HFD control and DSS treated mice (n= 4 mice per group). (**d**) Gene expression of *Cd36* in cecum tissue from LFD and HFD control and DSS treated mice (n= 5 LFD/HFD, n=7 LFD/HFD+DSS per group). (**e**) Flow cytometry analysis for surface expression of CD36 in intestinal macrophages in LFD and HFD control and DSS treated mice (n= 4 mice per group). (**f**) Representative immunofluorescence staining for milk-fat globule-EGF factor 8 (Mfge8, green), nuclei (DAPI, blue) and apoptotic cells (TUNEL, red) in LFD and HFD control and DSS treated mice (n= 5 mice per group). (**g**) Signal ratio of Mfge8 from HFD and LFD treated mice on day 9 post DSS per high-powered field (n=5 mice per group average of 3 images per mouse). Data are presented as mean ± SEM. Statistical significance was analyzed by Student t Test (**a, b, d, e**).

To confirm reduced efferocytosis in vivo, we examined downstream signaling pathways induced by clearance of apoptotic cells in sorted macrophages. After efferocytosis, phospholipids and cholesterol derived from the membranes of apoptotic cells need to be efficiently transported out of macrophages to protect against macrophage oxidative stress and cell death^31^. Macrophage lipid homeostasis is a tightly regulated process and imbalance can lead to altered macrophage functions that can promote inflammatory diseases such as atherosclerosis^32^. Cholesterol efflux receptors ABCA1 and ABCG1 play critical roles in this process, transporting phospholipids and cholesterol out of cells. Recognition of apoptotic cells induces upregulation of ABCA1 and ABCG1^33^ with lack of ABCA1 resulting in reduced efferocytosis^34^. We assessed ABCA1 and ABCG1 expression after DSS treatment and find macrophages from HFD fed mice have lower ABCA1 expression as compared to macrophages from LFD fed mice (Fig. 5b). Expression of ABCG1 was unchanged in both groups (data not shown). The lack of ABCA1 upregulation indicates macrophages from HFD mice have not taken up and/or processed apoptotic cells.

Efferocytosis is induced by macrophage recognition of phosphatidylserine (PS). PS is normally confined to the cytoplasmic side of the plasma membrane. In cells undergoing apoptosis, externalized PS is an “eat me” signal resulting in cell uptake^35^. Macrophages express multiple receptors that recognize externalized PS, including Axl, Mertk^36^, integrins αVβ3 and αVβ5 and their co-receptor CD36^37^. Loss of efferocytosis receptors Axl and Mertk or CD36 increase sensitivity to intestinal damage with decreased macrophage uptake of apoptotic cells^38-40^. Macrophages deficient for CD36 also have defective IL-10 expression after exposure to apoptotic cells^39,40^. While we find expression of Axl, Mertk, and αVβ3/5 were not altered by HFD or after DSS treatment (Fig. 5c and Supplementary Fig. 14a-c), we find reduced CD36 transcription in intestinal tissue (Fig. 5d), as well as a decreased surface CD36 expression by intestinal macrophages (Fig. 5e).

Uptake of apoptotic cells through αVβ3/αVβ5/CD36 requires bridging protein milk fat globule-EGF factor-8 (Mfge8)^37,41^. In mouse models of colitis, Mfge8 is constitutively expressed, with increased expression after intestinal damage that returns to baseline after tissue healing^42^. Initially, we detected increased tissue Mfge8 in both LFD and HFD treated mice. However, while Mfge8 decreased in LFD mice after damage resolution, levels remained elevated in HFD mice (Fig. 5f, g). These findings demonstrate that mechanisms to clear apoptotic cellular debris are induced in HFD after tissue injury but are not effective in promoting apoptotic cell clearance and tissue repair. Together, these data suggest that dietary lipids interfere with CD36-mediated recognition and internalization of apoptotic neutrophils.

### HFD impairs macrophage efferocytosis by blocking of CD36

To test whether lipid exposure directly interfered with macrophage uptake of apoptotic cells, we pretreated BMDMs with oleic acid. We then incubated the BMDMs with apoptotic neutrophils and assessed efferocytosis. After pretreatment with oleic acid, we found BMDMs had decreased capacity to engulf apoptotic neutrophils with a decreased proportion containing a single neutrophil, and none containing more than one cell (Fig. 6a-c). Lipid pre-treatment of apoptotic neutrophils prior to culturing with BMDMs did not impact BMDM uptake of neutrophils (Supplementary Fig. 15 and data not shown). The first step in efferocytosis is tethering the apoptotic cell to the macrophage. We found that pre-treatment with oleic acid decreased ability of BMDMs to tether apoptotic neutrophils (Fig. 6d). This lack of tethering likely underlies the reduced efferocytosis we observe.

**Figure 6.**
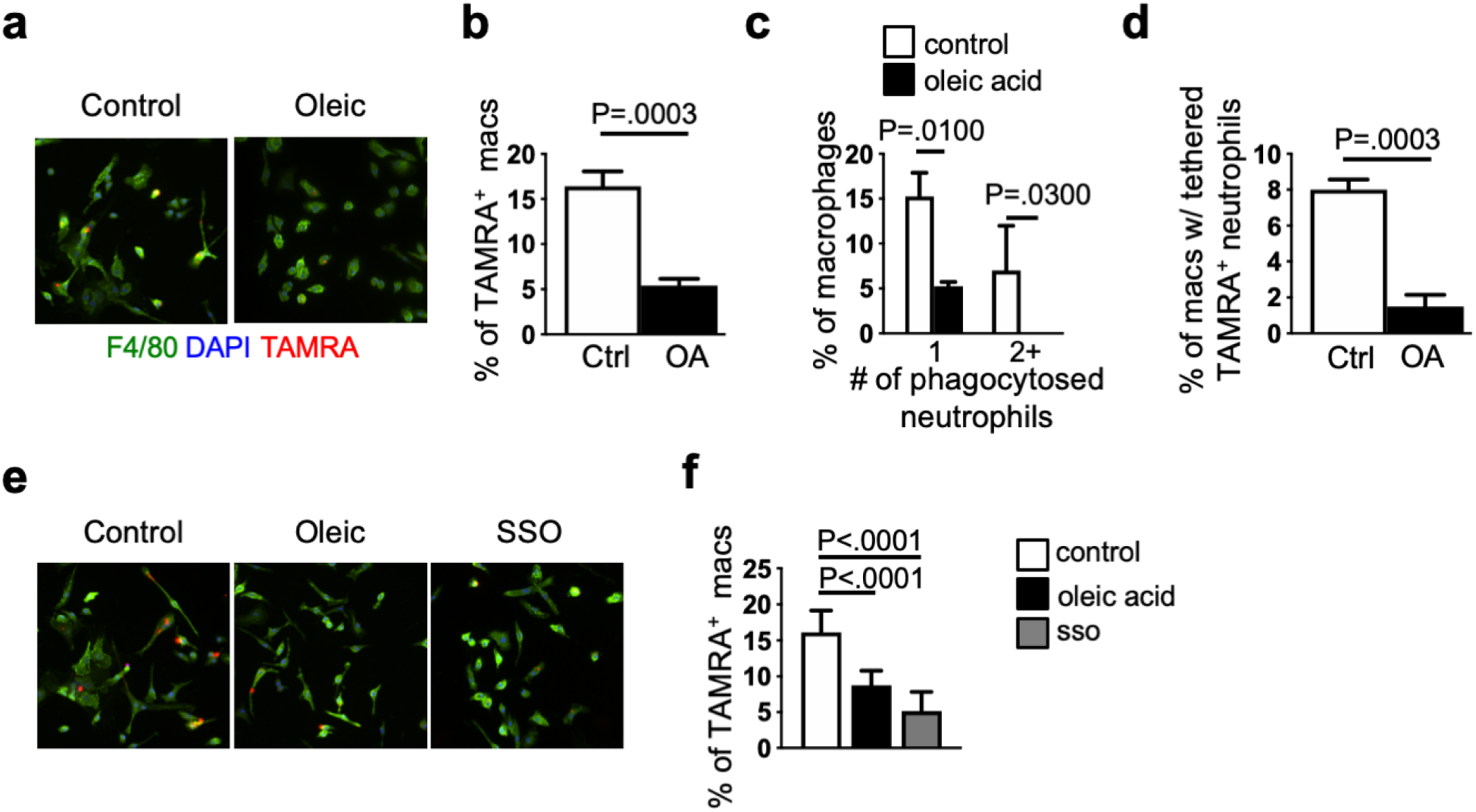
Lipids directly impair macrophage efferocytosis through blocking of CD36. (**a**) Representative immunofluorescence image from phagocytosis assay (stained for macrophages (F4/80, green), nuclei (DAPI, blue) and apoptotic neutrophils (TAMRA, red) in control and oleic acid treated bone marrow derived macrophages. (**b**) Quantification of phagocytic capacity. The percent of TAMRA^+^ macrophages is indicative of the percentage of macrophages that phagocytosed TAMRA^+^ apoptotic neutrophils (n=7 replicate experiments). (**c**) Quantification of the percent of macrophages that have phagocytosed 1 or 2 or more TAMRA+ neutrophils after control and oleic acid treatment. (**d**) Quantification of percent of macrophages with tethered TAMRA positive neutrophils after control or oleic acid pretreatment (control (n=7), oleic acid (n=8)) in (**a**). (**e**) Representative immunofluorescence image from phagocytosis assay (stained for macrophages (F4/80, green), nuclei (DAPI, blue) and apoptotic neutrophils (TAMRA, red) in control, oleic acid, and SSO treated bone marrow derived macrophages. (**f**) Quantification of phagocytic capacity as (**e**) in control, oleic acid and SSO treated BMDMs (n= 4 experiments). Scale bar equals 100 microns. Data are presented as mean ± SEM. Statistical significance was analyzed or Student’s T test (**b, d**,) or One-way Anova with Tukey’s posttest (**c, f**).

In addition to its function as an efferocytosis receptor, CD36 is also a receptor for dietary and endogenous lipids^43^. To assess whether lipid exposure directly induced changes in CD36 expression, we pre-treated BMDMs with oleic acid and found decreased staining with CD36 antibodies (Supplementary Fig. 16). It was not clear if this was inducing loss of CD36 from the cell surface or blocking interactions between CD36 and anti-CD36 antibodies or apoptotic cells. To resolve this question, we pretreated BMDMs with Sulfo-N-succinimidyl Oleate (SSO), a lipid compound that irreversibly binds to CD36 but cannot be internalized^44^. As with oleic acid pretreatment, SSO pretreatment blocked antibody binding of CD36 (Supplementary Fig. 16). We found that SSO treatment decreased uptake of apoptotic neutrophils in BMDMs as compared to control treated BMDMs (Fig. 6e, f). These findings demonstrate that dietary lipids interfere with CD36 accessibility, limiting macrophage efferocytosis of apoptotic neutrophils.

## Discussion

Our results indicate that high fat diets not only increase systemic inflammation, but also directly interfere with intestinal barrier repair by disrupting interactions between apoptotic neutrophils and CD36, an efferocytosis receptor. Efferocytosis turns on key effectors for barrier repair, including IL-10, which signals through epithelial cells to reestablish gap junctions. Efferocytosis is also a key signal to down regulate chemokines that recruit neutrophils into tissue^45^. Together, this results in increased tissue accumulation of apoptotic cells which can further amplify tissue inflammation and impair barrier repair. Lipids and apoptotic cells share a receptor, CD36 and we find dietary lipids can interfere with apoptotic cell binding to CD36. In this way, excess lipids antagonize key anti-inflammatory and barrier repair signals, including macrophage IL-10 secretion, that are required to reestablish intestinal integrity and limit microbial penetration into the tissue. Together, microbial signals and chemokines continue to recruit neutrophils into the tissue, further potentiating tissue damage and limiting tissue repair.

Recent data have demonstrated that, in the steady state, intestinal macrophages and dendritic cells uptake of apoptotic epithelial cells induces a homeostatic transcriptional program that promotes regulatory T cells^46^. It remains to be determined if the same PS receptors are utilized in this homeostatic pathway and further if dietary lipids also interfere in apoptotic cell clearance at steady state. Defects in similar clearance pathways are observed in several inflammatory disorders and it will be important to understand if dietary lipids interfere with apoptotic neutrophil clearance in non-intestinal sites. As the PS receptors, Axl and Mertk have also been shown to be important for barrier repair^38,47^, understanding the context dependent cues that bias towards utilization of one receptor over another will allow for development of therapeutic interventions to increase uptake of apoptotic cells to improve barrier repair and resolve inflammatory conditions such as inflammatory bowel disease.

## Methods

### Mice

Male C57BL/6J (JAX #000664), CX_3_CR1^+^ GFP/+ (JAX # 005582), CX_3_CR1-CreERT2 (JAX# 021160) and Lgr5-EGFP-IRES-creERT2 (JAX# 008875) were purchased from Jackson Labs. L-10 receptor α conditional (IL-10Rα ^fl/fl^) mice were from Dr. Werner Muller^48,49^. All mice were kept and bred under specific pathogen-free (SPF) conditions at the animal facility of Baylor College of Medicine. All mouse experiments were performed with at least 4 mice per group in male mice between 6 and 8 weeks of age. Multiple experiments were combined to assess statistical significance. Littermate controls were used for each experiment and mice were randomly assigned to experimental groups. All experiments were performed in accordance with approved protocols by the Institutional Animal Care and Usage Committee at Baylor College of Medicine.

### Short term diet feeding and intestinal injury

Mice were fed 10% kcal low fat diet (LFD) (Research Diets, D12450B) or 60% kcal high fat diet (HFD) (Research Diets, D12492) ad libitum for one week prior to treatment with: 1) 2% Dextran sodium sulfate (DSS, ThermoFisher, AAJ1448922) placed in drinking water for 5 days followed by plain drinking water. 2) a single intraperitoneally (i.p.) injection of 20mg/Kg BW of doxorubicin (Sigma Aldrich, D1515). 3) C. rodentium DBS100 (ATCC 51459; American Type Culture Collection) was harvested at log phase growth and 10^10^ CFU in saline were delivered by oral gavage.

### Glucose tolerance and Insulin tolerance test

GTT and ITT were performed by the Mouse Metabolic Phenotyping Center at Baylor College. Studies were performed in mice fed LFD and HFD for two and eight weeks. ***Glucose tolerance test (GTT):*** After a 6h overnight fast, 1.5g/Kg body weight of glucose was given i.p. to each mouse. Blood was collected from tail vein at 0, 15, 30, 60 and 120 min and glucose levels were checked using a glucometer (Life Scan, Milpitas, CA). ***Insulin tolerance test (ITT):*** For ITT, 1 U/kg body weight insulin (HUMULIN R) was injected intraperitoneally to mice after a 4h fast. Blood glucose was measured as described above.

### Histology

Cecum was fixed in Carnoy’s fixation for 1-2 days before being placed in methanol prior to paraffin embedding. 4µM sections were deparaffinized and stained with hematoxylin or Alcian Blue/PAS. Images were taken with a Nikon Ti Eclipse microscope. Sections from 5 mice were used for blinded colitis scoring according to established criteria^29,50^. The number of Alcian blue/PAS positive goblet cells were counted per villi for 10 villi in 5 mice per group.

### Immunofluorescence tissue staining

Sections were fixed, embedded and deparaffinized as described above. Sections were permeabilized, blocked, and stained overnight at 4°C with the following primary antibodies at a 1:100 dilution: Ki67 (Novus Biologicals, NB110-89719), MUC2 (Cloud Clone, PAA705Mu02), anti-Ly6g 1A8 (BioCell Cat# BE0075-1), F4/80 (Abcam, ab6640), CD11b (Abcam, ab184307), anti-MFGE8 (MBL, D199-3). Sections were washed, stained with the following secondary antibodies at a 1:200 dilution at room temperature for 1h: anti-rabbit NorthernLights-557 (R&D systems, NL007), anti-rat Alexa 488 (Cell signaling, #4416), anti-rabbit NorthernLights-637 (R&D systems, NL005), anti-hamster 488 (Abcam, ab173003). Followed by DAPI (Sigma Aldrich) and mounted using Aqua Mount (Polysciences) anti-fade mounting media and coverslipped. In Situ Cell Death Detection Kit TMR red (TUNEL) (Sigma/Roche, 12156792910) staining was performed according to manufactures instructions prior to immunofluorescence staining. Images were taken on Nikon Ti Eclipse microscope. Mice were injected with BRDU (BD Pharmingen, 550891) 2h before collection and fixation of tissue. Signal ratio for MFGE8 was assessed using Image J software where signal ratio was determined by dividing the specific fluorescence intensity of MFGE8 by the overall general fluorescence (background) of the image.

### DNA preparation and 16S (v4) rRNA gene Sequencing

Fecal DNA extraction was performed using the PowerMag Microbiome DNA isolation kit (MoBio) following manufacturer’s instructions. Sequencing of the 16S rRNA V4 region were performed as described^51^.

### Bone-marrow derived macrophages (BMDMs)

BMDMs were differentiated from 8-week-old male and female C57BL/6 mice as previously described^52^. Single cell suspension of bone marrow cells was cultured for 6 days in 50% DMEM (Corning) supplemented with, 20% FBS, 30% L cell (ATCC CRL-2648) media, 2mM glutamine, 1 mM pyruvate, 1 unit/ml pen/strep, and 55μM β-ME. Confirmation of macrophage differentiation was assessed by flow cytometry as described below. All assays were performed in DMEM supplemented with 10% FBS, 1 unit/ml pen/strep and 1mM HEPES.

### Apoptotic neutrophils

Neutrophils were isolated from bone marrow using a density gradient (Histopaque 1077 and 1099) as previously described^53^ and incubated for 24 hours at 37°C in DMEM containing 1% FBS. Apoptosis was assessed by trypan blue staining. For phagocytosis assays, apoptotic neutrophils were stained with TAMRA (ThermoFisher, C1171) according to manufactures instructions.

### Gene expression

RNA from whole cecum or sorted macrophages was isolated using Trizol (Invitrogen) according to manufacturer’s protocol. cDNA was synthesized using iScript reverse transcription kit (Bio-rad Laboratories). Real-time quantitative qPCR was performed using SYBR Green Supermix (Bio-rad Laboratories) using a CFX384 Touch real-time PCR machine. Thermocycling program was 95°C for 2 min followed by 40 cycles at 95°C for 15 s, 60°C for 30s, and 72°C for 30s. The following primers were used: mIL-10-F: CCAGCTGGACAACATACTGCT, mIL10-R: CATCATGTATGCTTCTATGCAG, mCD36-F: CCGAGGACCACACTGTGTC, mCD36-R: AACCCCACAAGAGTTCTTTCAAA, mGAPDH -F: AATGTGTCCGTCGTGGATCT, mGAPDH: CATCGAAGGTGGAAGAGTGG, mABCA1-F: GGACATGCACAAGGTCCTGA, mTBP-F: ACCGTGAATCTTGGCTGTAAAC, mTBP-R: GCAGCAAATCGCTTGGGATTA, mABCA1-R: AGAAAATCCTGGAGCTTCAAA, mAxl-F: GTTTGGAGCTGTGATGGAAGGC, mAxl-R: CGCTTCACTCAGGAAATCCTCC, mMerTk-F: CAGGAAGATGGGACCTCTCTGA, mMerTk-R: GGCTGAAGTCTTTCATGCACGC, mTNF-F: TGGGAGTAGACAAGGTACAACCC, mTNF-R: CATCTTCTCAAAATTCGAGTGACA. mOccludin-F: TCAGGGAATATCCACCTATCACCTCAG, mOccludin-R: CATCAGCAGCAGCCATGTACTCTTCAC, mZO-1-F: AGGACACCAAAGCATGTGAG, mZO-1-F: GGC ATTCCTGCTGGTTACA, mCXCL1-F: TGAGCTGCGCTGTCAGTGCCT, mCXCL1-R: AGAAGCCAGCGTTCACCAGA, mCXCL2-F: GAGCTTGAGTGTGACGCCCCCAGG, mCXCL2-R: GTTAGCCTTGCCTTTGTTCAGTATC. Relative expression of target gene was determined using the delta delta CT method. GAPDH and TBP were used as internal controls.

### Phagocytosis assay

Sorted intestinal macrophages isolated from LFD and HFD mice were incubated at a 1:2 ratio with TAMRA loaded apoptotic neutrophils for 1h in FACS tubes and cytospun before immunostaining. BMDMs were plated at 1X10^6^ cells in cover glass MakTek dish (MakTek Corporation) or 1.5X10^6^ per well in 24 well TC plates and incubated at a 1:3 ratio with TAMRA loaded apoptotic neutrophils for 1h before RNA isolation or immunostaining. Oleic acid pre-treated apoptotic neutrophils were used as an additional control where indicated.

### Lipid Treatment of BMDMs

Fatty acids were dissolved in ethanol as described^6^. BMDMs were treated with 400µm oleic acid (Nu-Chek Prep)^6^ or treated with 2µg/ml of Sulfo-N-succinimidyl Oleate (SSO) (Sigma, SML2148) or equivalent amount of solvent (ethanol). Viability was determined using Presto Blue (ThermoFisher, A13261) assay according to manufactures instructions.

### Neutrophil depletion

Mice were then injected once i.p. with 400µg of IgG2a isotype control (BioCell Cat# BE0089) or anti-Ly6g (BioCell Cat# BE0075-1).

### Lamina propria cell isolation

Isolation of lamina propria cells were performed as previously described^29,54^. In short, large intestinal tissue was placed in PBS, cut open to expose the lumen and luminal contents were removed. Intestine were cut in 1 cm section and then treated with 1mM DTT and 30mM EDTA for 10min to remove mucus and epithelial layer. Tissue was then digested in collagenase 8 (Sigma-Aldrich) and DNase-containing media supplemented with 10% FBS while shaking at 37 degrees for 1hr followed by separation on a 40%/80% Percoll (Sigma Aldrich) gradient.

### Flow cytometry and FACS sorting

Flow cytometry and analysis were performed with an LSR II (BD) and FlowJo software (Tree Star). Dead cells were excluded using the Live/Dead fixable aqua dead cell stain kit (Invitrogen). Macrophage populations were sorted on a FACSAria Cell sorter (BD Biosciences). Pooled samples from 5-7 mice were used to obtain an n=1 for a total “n” equaling 4-5 for each group for gene expression and statistical analysis. The following antibodies were used for flow staining and or sorting: MHC II (M5/114.15.2, BioLegend Cat# 107620), CD11b (M1/70, Biolegend Cat# 101226), CD11c (N418, Biolegend, Cat # 117317), Ly6C (1A8, BD PharMingen Cat#560525), CD36 (HM36, Biolegend CA# 102605), αvβ5 (ALULA, BD PharMingen Cat#565836), αvβ3 (RMV-7, BD PharMingen Cat#104106), CD45 (30-F11, Biolegend Cat#103149), CD103 (2E7, BioLegend Cat#121413), DAPI (Sigma-Aldrich Cat# D9542).

### Oil-red-O staining

Sorted macrophages were seeded on microscopy slides for 2 h and then fixed in 4% PFA for 15 min and washed twice with distilled water. Slides were placed in 100% propylene glycol for 5 min then washed twice with distilled water. Slides were then stained with Oil-red-O for 15min at 65°C. After staining, slides were washed in 85% propylene glycol for 5min then washed 2X with distilled water. Slides were then counterstained with hematoxylin for 30s and washed thoroughly with tap water and mounted using aqueous mounting media. Images were taken using a Leica brightfield microscope. BMDMs treaded with overnight with oleic acid were used as a positive control for staining.

### Overexpression of IL-10

Plasmid DNA expression of control or IL-10 (InVivoGen) were delivered intravenously (i.v.) at 10µg DNA/mouse diluted in TransIT-EE Hydrodynamic Delivery solution (Mirus) at 0.1 ml/g body weight on 1 day after start of DSS treatment.

### 4-hydroxy tamoxifen (4OHT) administration

(Z)-4-Hydroxytamoxifen, 98% Z isomer (4OHT, Sigma) was resuspended to 20mg/ml in ETOH with heating to 37°C. 4OHT was diluted in corn oil (Sigma) and mice were injected I.P. with 0.2mg every 3 days (CX_3_CR1CreERT2 and control) starting on Day 0 of DSS treatment or on 2 sequential days one week before the start of DSS treatment (LGR5-CreERT2 and control).

### Statistical analysis

One-way analysis of variance (ANOVA) with Tukey’s posttest or unpaired t test was performed using a 95% confidence interval. All analyses were performed using GraphPad Prism version 8.0. Differences were considered to be significant at P values of < 0.05.

### Contributions

G.E.D. and A.A.H. designed experiments and wrote the manuscript with the input from all co-authors. A.A.H. and M.K. performed, designed and analyzed the experiments. D.F.Z.R., A.M.J., H.W.S., M.C.R., W.-J.W. and K.N., performed experiments.

## Acknowledgements

This work was supported by NIH AI125264 (G.E.D.), institutional NRSA T32AI053831_Corry (A.A.H.), AAI Careers in Immunology Fellowship (M.K.), the Texas Medical Center Digestive Disease Center Cellular and Molecular Morphology Core and Pilot Fund (G.E.D) (NIH P30DK056338), the Cytometry and Cell Sorting Core at Baylor College of Medicine with expert assistance of Joel M. Sederstrom and funding from the NIH (P30 AI036211, P30 CA125123, and S10 RR024574), Center for Metagenomics and Microbiome Research for expertise in microbiome analysis and Buck Samuels for use of microscope.

