## Supplementary Figures for "Defective intestinal repair after short-term high fat diet due to loss of efferocytosis"

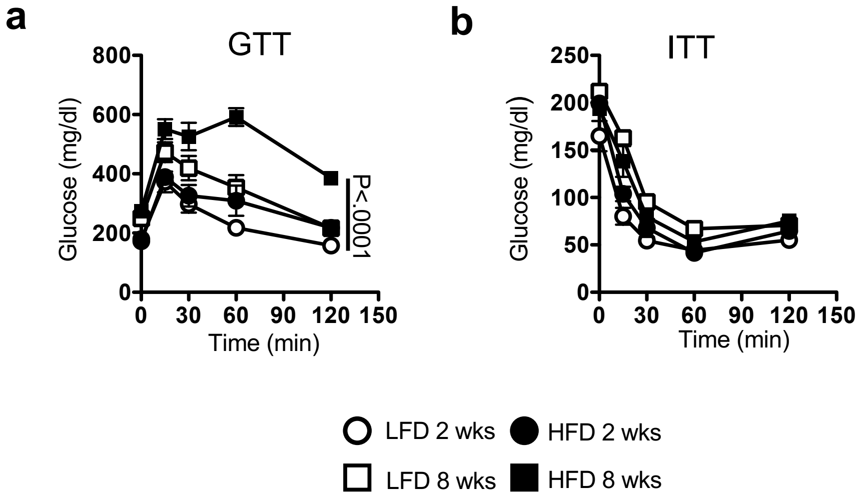


**Supplementary Figure 1**. **Short-term diet feeding does not lead to major metabolic changes.** Glucose tolerance test (**a**) and insulin tolerance test (**b**) of mice treated with LFD or HFD for 2 or 8 weeks. Data are presented as mean ± SEM, 10 mice per group. Statistical significance was analyzed by Student t Test.

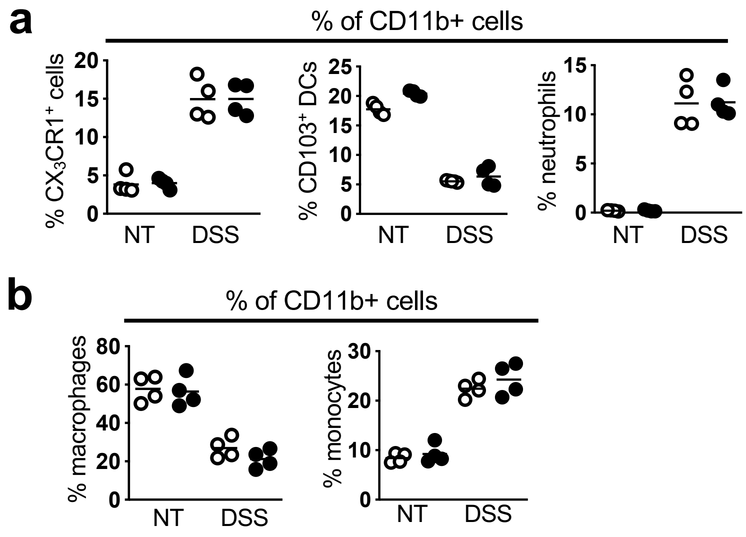


**Supplementary Figure 2**. **Intestinal myeloid cell population are similar in LFD and HFD mice before at day 5 of DSS challenge.** (**a**) Flow cytometry analysis of $CX_3CR1$+, CD103+, and ly6g+ cell frequencies from total CD11b+ cells in intestine of LFD and HFD exposed mice before and at day 5 of DSS treatment. (**b**) Macrophage and monocytes frequencies from total $CX_3CR1$+ CD11b+ cells in (**a**). Data are presented as mean ± SEM, 4 mice per group.

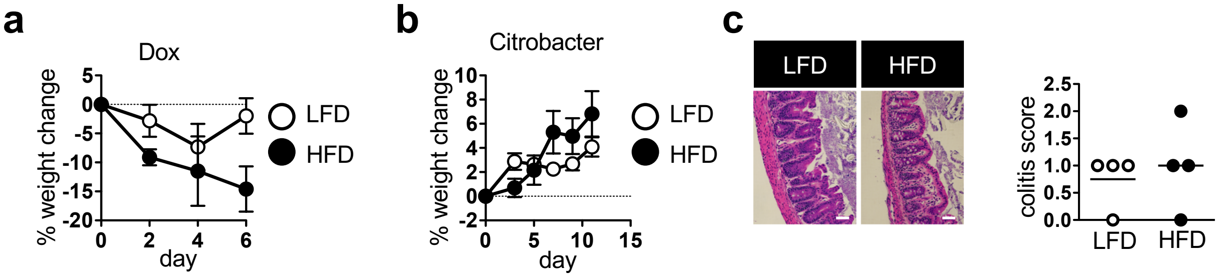


**Supplementary Figure 3. HFD feeding sensitizes to chemotherapy but not infectious colitis.** (**a**) Body weight change of LFD and HFD treated mice injected with 20mg/kg BW of doxorubicin (n=12 mice per group). (**b**) Body weight of LFD and HFD mice infected orally with $10^{10}$ CFU C. *rodentium* (n=5 mice per group). (**c**) Representative H&E staining in colon and blinded colitis scores on day 11 post infection (n=5 mice per group). Data are presented as mean ± SEM. Scale bar equals 100 microns.

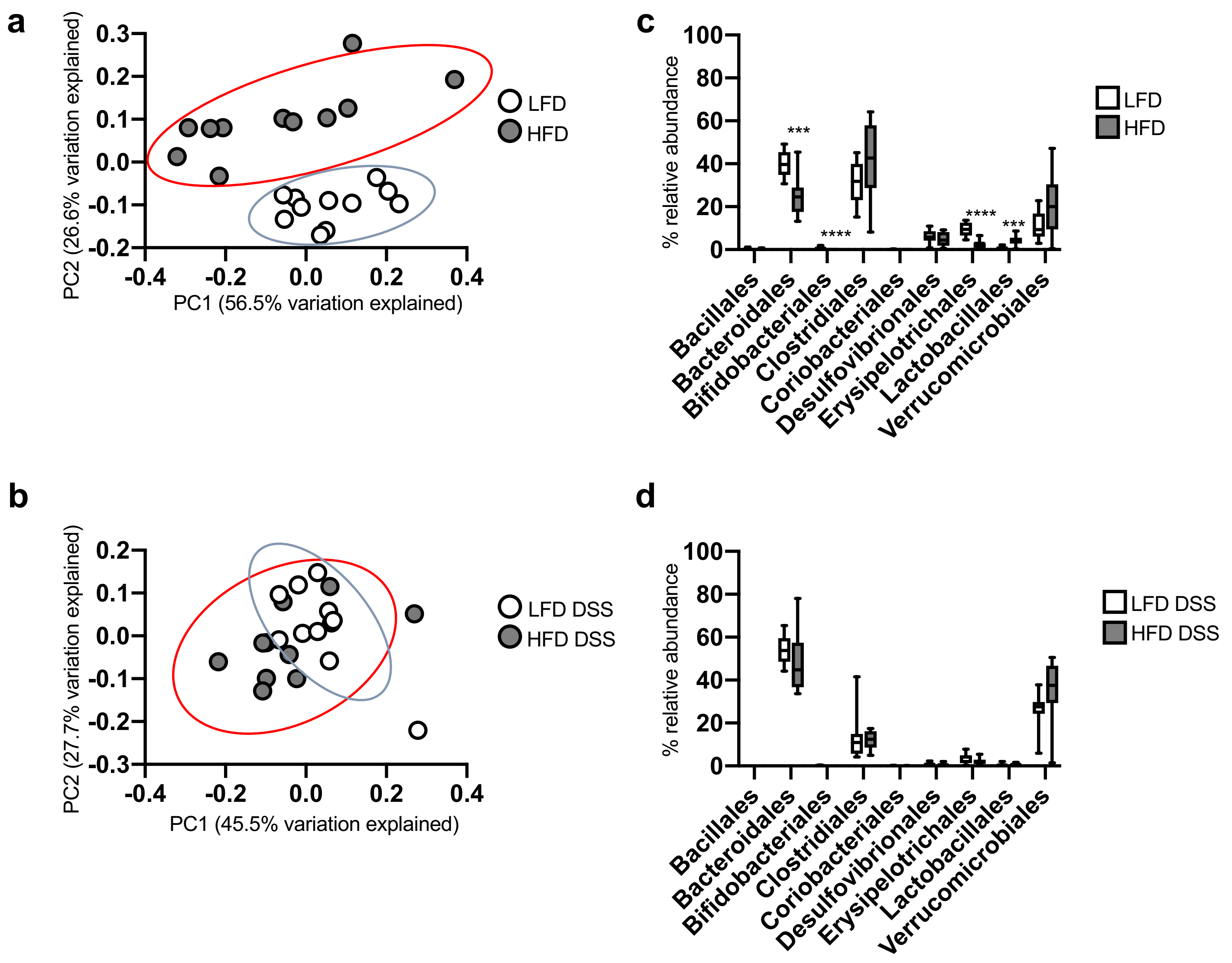


**Supplementary Figure 4. Microbiota changes after 1 week of HFD treatment are lost after DSS challenge.** (**a-b**) Principal coordinate analysis (PCoA) of weighted UniFrac clustering of Beta Diversity of the gut microbiota in LFD and HFD treated mice before and after DSS treatment as measured by 16s rRNA analysis. Each dot represents a single mouse. (**c-d**) Relative abundance at Taxa level shown. Data are presented as mean ± SEM, n=11 samples per group. Statistical significance was analyzed by Student t Test (**b**). ***p< 0.001, p<0.0001****.

**a**

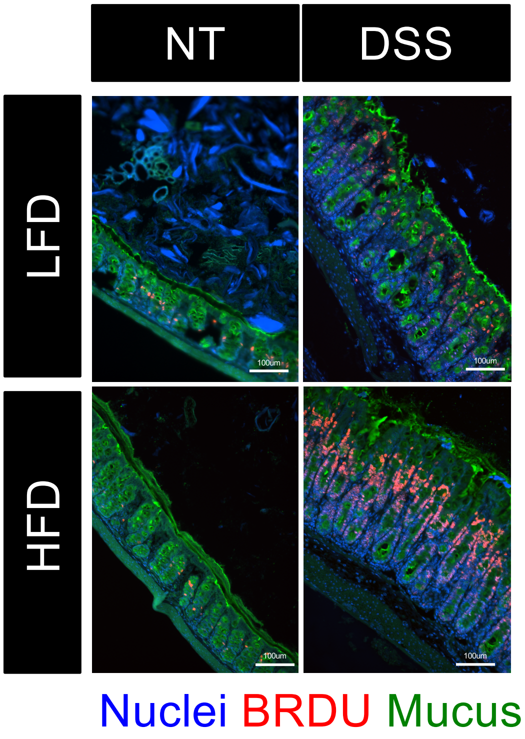


**Supplementary Figure 5. Increased proliferation in intestinal tissue of HFD treated mice at late timepoints after DSS challenge.** (**a**) Representative immunofluorescence staining for Muc2 (mucus, green), nuclei (Dapi, blue) and BRDU (proliferation, red) in LFD and HFD fed DSS treated mice. Scale bar equals 100 microns.

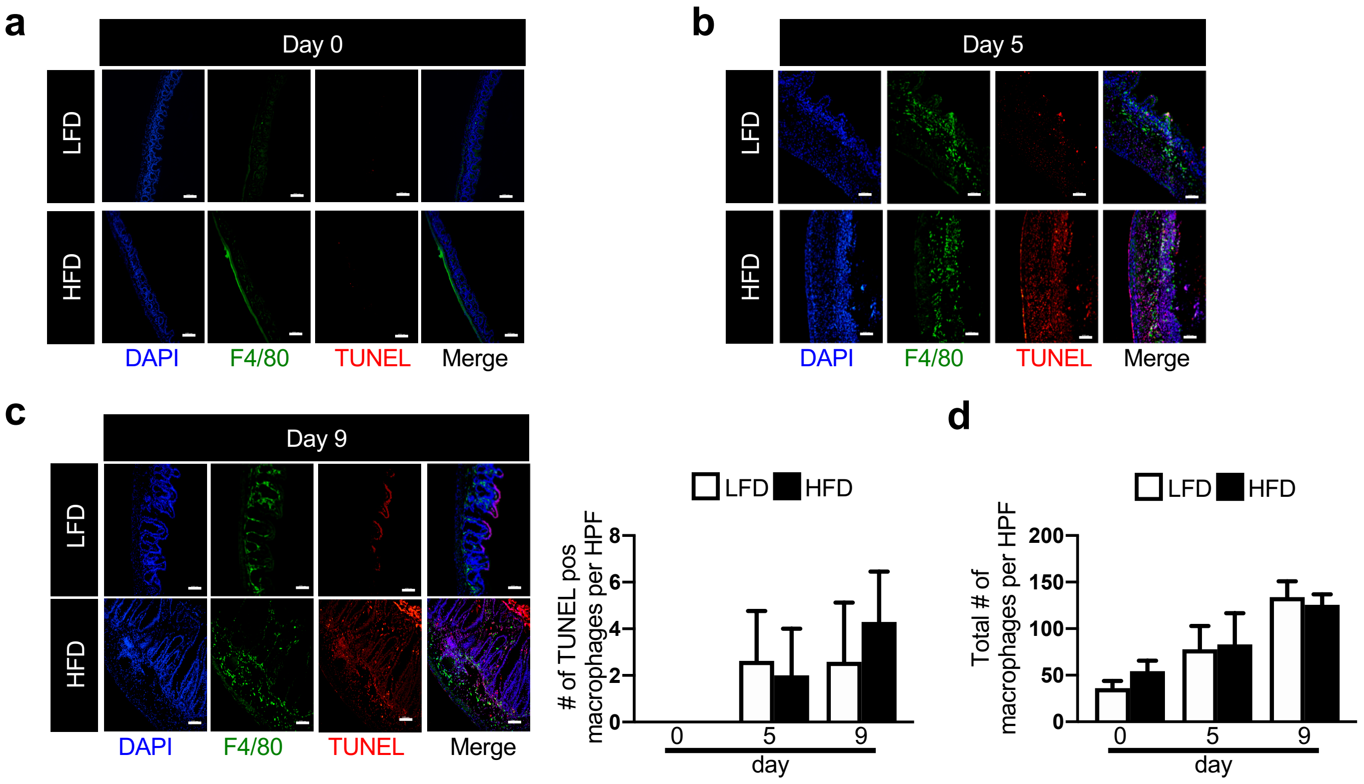


**Supplementary Figure 6. Tissue macrophages increase after DSS challenge.** (**a-c**) Representative immunofluorescence staining for F4/80 (green), nuclei (Dapi, blue), apoptotic cells (TUNEL, red), CD11b (magenta) in LFD and HFD DSS treated mice and day 0 and 5 after DSS treatment. Scale bar equals 100 microns. Total (**d**) and apoptotic (**c**) macrophage numbers in LFD and HFD treated mice before and at day 5 and 9 after DSS treatment. n=4 per group.

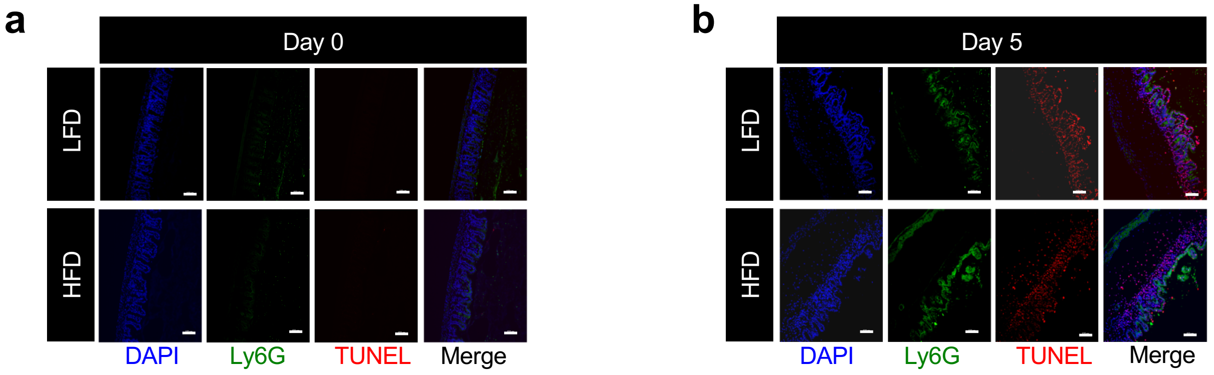


**Supplementary Figure 7. Apoptotic neutrophil numbers are similar in intestinal tissue of HFD treated mice at day 5 of DSS challenge.** (**a-b**) Representative immunofluorescence staining for Ly6G (green), nuclei (Dapi, blue), apoptotic cells (TUNEL, red), CD11b (magenta) in LFD and HFD DSS treated mice and day 0 and 5 after DSS treatment. Scale bar equals 100 microns.

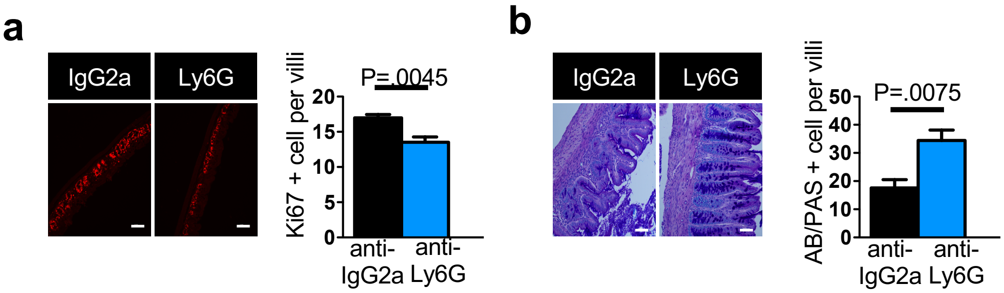


**Supplementary Figure 8. Neutrophil depletion reduces intestinal cell proliferation and restore goblet cells in HFD treated mice after DSS treatment.** (**a**) Representative immunofluorescence staining for proliferating cells (Ki67, red) and quantification of Ki67 positive cells from mice in Figure 2 (n=5 mice per group). (**b**) Representative Alcian Blue/PAS staining and quantification of goblet cells in cecum of mice in Figure 2 (n= 5 mice per group). Data are presented as mean ± SEM. Statistical significance was analyzed by Student t Test. Scale bar equals 100 microns.

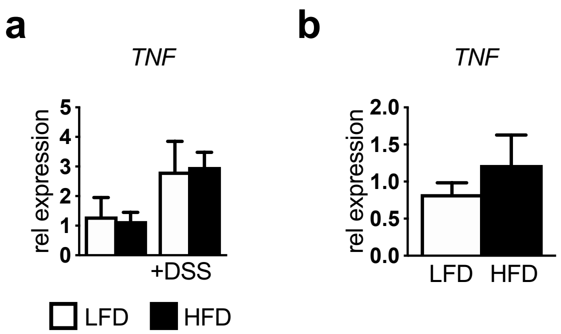


**Supplementary Figure 9. Short term HFD feeding does not increase macrophage inflammatory cytokines**. (**a**) Cecal gene expression of TNFα in LFD and HFD control and DSS treated mice (n=6 LFD, n=6 HFD, n= 7 LFD DSS, 10=HFD DSS mice per group). (**b**) Gene expression of TNFα in sorted intestinal macrophages from LFD and HFD exposed DSS treated mice. Data are presented as mean ± SEM, n=5 mice per group for cecum gene expression. n= 5 per group for sorted cells gene expression where sorted cells from 5 mice pooled per "n" per group.

**a**

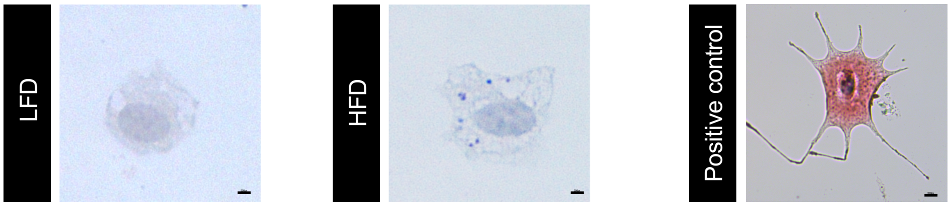


**Supplementary Figure 10. Short-term diet feeding does not induce lipid accumulation in intestinal macrophages.** (**a**) Oil-red-O staining of sorted intestinal macrophages from LFD and HFD fed mice. Positive control is oleic acid pretreated bone-marrow derived macrophage. Scale bar equals 500px.

**a**

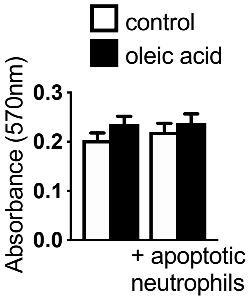


**Supplementary Figure 11. Oleic acid treatment does not induce macrophage cell death.** (**a**) Presto Blue cell viability assay in control or oleic acid treated BMDMs alone or after exposure to apoptotic neutrophils (n= 3 experiments). Data are presented as mean ± SEM.

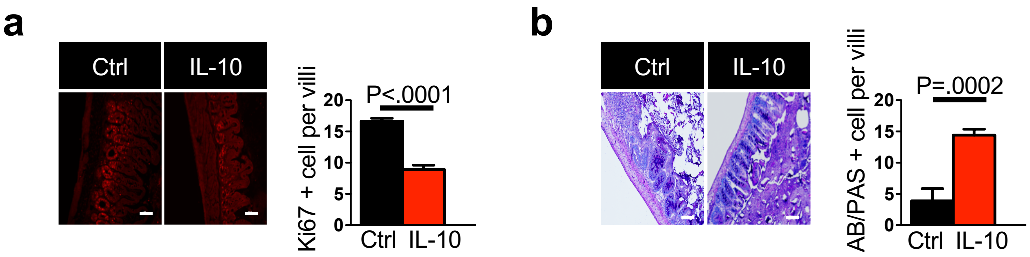


**Supplementary Figure 12. Overexpression of IL-10 reduces intestinal cell proliferation and restore goblets cell in HFD treated mice after DSS.** (**a**) Representative immunofluorescence staining and quantification of proliferating cells (Ki67, red) of mice in Figure 3. (**b**) Representative Alcian Blue/PAS staining and quantification for goblet cells in cecum of mice in Figure 3. Data are presented as mean ± SEM, 5-6 per group imaging samples and quantification. Statistical significance was analyzed by Student t test. Scale bar equals 100 microns.

**a**

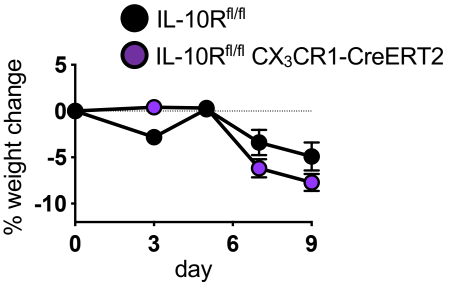


**Supplementary Figure 13. IL-10R on macrophages is not required for IL-10 mediated protection for intestinal damage in HFD.** (**a**) Weightloss in tamoxifen treated IL-10R$^{fl/fl}$ (control) and IL-10R$^{fl/fl}$ CX$_3$CR1-CreERT2 mice treated with HFD and DSS and hydrodynamic delivery of an IL-10–producing plasmid (n= 5 per group).

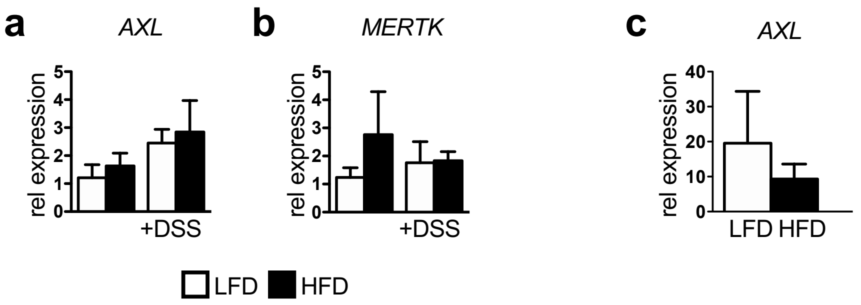


**Supplementary Figure 14. No change in AXL or MERTK expression after HFD.** (**a-b**) Cecum gene expression of AXL and MERTK in LFD and HFD control and DSS treated mice (n= 5 mice per group). (**c**) Gene expression of AXL in sorted intestinal macrophages from LFD and HFD DSS treated mice. Data are presented as mean ± SEM, n= 5 per group for sorted cells gene expression where sorted cells from 5 mice pooled per “n” per group. n=5 mice per group for cecum gene expression.

**a**

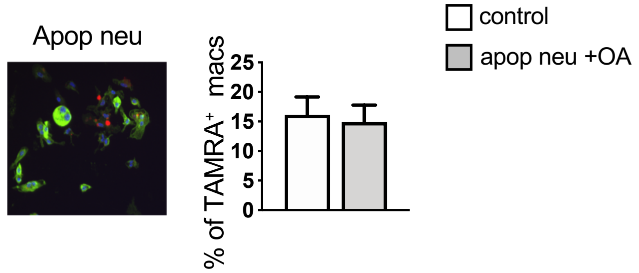


**Supplementary Figure 15. Lipid pretreatment of apoptotic neutrophils does not impair macrophage efferocytosis.** (**a**) Immunofluorescence staining and quantification of percentage of TAMRA+ macrophages after treatment with control or lipid-treated apoptotic neutrophils.

**a**

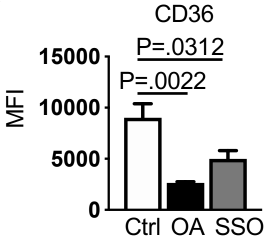


**Supplementary Figure 16. Oleic acid and SSO inhibit antibody recognition of CD36.** (**a**) Mean fluorescence intensity of CD36 in control, oleic acid or SSO treated BMDMs. (n= 4 experiments). Data are presented as mean ± SEM.
